# Making plants into cost-effective bioreactors for highly active antimicrobial peptides

**DOI:** 10.1101/583146

**Authors:** Meron R Ghidey, S M Ashiqul Islam, Grace Pruett, Christopher Michel Kearney

**Affiliations:** Biomedical Studies Program, Baylor University, Waco, TX, USA; Department of Biology, Baylor University, Waco, TX, USA

**Keywords:** antimicrobial peptides, plant expression, *Nicotiana benthamiana*, elastin-like peptide, heterologous protein expression.

## Abstract

As antibiotic-resistant bacterial pathogens become an ever-increasing concern, antimicrobial peptides (AMPs) have grown increasingly attractive as alternatives. Potentially, plants could be used as cost-effective AMP bioreactors; however, reported heterologous AMP expression is much lower in plants compared to *E. coli* expression systems and often results in plant cytotoxicity, even for AMPs fused to carrier proteins. We wondered if there were a physical factor that made heterologous AMPs difficult to express in plants. Using a meta-analysis of protein databases, we determined that native plant AMPs were significantly less cationic than AMPs native to other taxa. To apply this finding to plant expression, we tested the transient expression of 10 different heterologous AMPs, ranging in charge from +7 to −5, in the the tobacco, *Nicotiana benthamiana*. We first tested several carrier proteins and were able to express AMPs only with elastin-like polypeptide (ELP). Conveniently, ELP fusion allows for a simple, cost-effective temperature shift purification. Using the ELP system, all five anionic AMPs expressed well, with two at unusually high levels (375 and 563 µg/gfw). Furthermore, antimicrobial activity against *Staphylococcus epidermidis* was an order of magnitude stronger (average MIC = 0.26 µM) than that typically seen for AMPs expressed in *E. coli* expression systems. Unexpectedly, this high level of antimicrobial activity was associated with the uncleaved fusion peptide. In contrast, all previous reports of AMPs expressed in both plant and *E. coli* expression systems show cleavage from the fusion partner to be required before activity is seen. In summary, we describe a means of expressing AMP fusions in plants in high yield, purified with a simple temperature-shift protocol, resulting in a fusion peptide with high antimicrobial activity, without the need for a peptide cleavage step.

## Introduction

The use of traditional antibiotics to control bacterial infections is threatened due to two undermining factors. First, drug discovery for new antimicrobial agents has been on the decline for the past three decades. The major classes of antibiotics have already been discovered and commercial incentives to develop new antibiotics have decreased (Charles and Grayson, 2004; Norrby, *et al.*, 2005; Schäberle and Hack, 2014). Second, the overuse of antibiotics has led to pathogenic and commensal bacteria incorporating and retaining genes for detoxification or export of antibiotics, inevitably resulting in resistance to all new antibiotics introduced (Aminov *et al*. 2010; Thung *et al.*, 2015; Enright *et al*. 2002; Nathan and Cars, 2014).

Both of these undermining factors are addressed by antimicrobial peptides (AMPs). First, the resources available to develop new AMP drugs is vast and recombinant peptide variants can be quickly generated, unlike the slow discovery and development cycle for antibiotics. AMPs are abundant across the taxa, being found in vertebrates, insects, fungi and plants. Thousands of AMPs have been isolated and tested experimentally (Wang *et al.*, 2015) and many more can be discovered using algorithms to scan genome data bases (Islam *et al.*, 2018b). Second, though resistance to AMPs has been shown to develop in bacteria (Kubicek-Sutherland *et al*., 2017), the multiple antimicrobial activities and low affinity targets typical of AMPs have been thought to make them more difficult targets for resistance development by pathogenic bacteria (Peschel and Sahl, 2006). From an environmental perspective, AMPs are not long-lasting in waste water, whereas low concentrations of antibiotics can induce resistance in soil and water-borne microbial communities (Bengtsson-Palme et al., 2018).

AMPs are not capable of completely replacing antibiotics, but could serve as replacements for some applications if they were produced at low cost. Though AMPs have been used clinically (Marr *et al.*, 2006), AMPs have a special potential for large-scale applications. Examples might include their use as a food preservative, as a topical disinfectant, or as a feed supplement for livestock or poultry. These sorts of applications would be dependent upon developing scalable and simple protocols for both production and purification.

Currently, there remain some roadblocks to developing these simplified protocols for large scale production. *E. coli* expression systems have been extensively demonstrated to effectively produce AMPs, but the AMP must be fused to a carrier protein in order to protect the bacterium from antimicrobial activity (Li, 2009). Various fusion partners have been used, such as SUMO (Li *et al.*, 2009a; Zhang *et al.*, 2015), GST (Liang *et al.*, 2006) and TRX (Tian *et al.*, 2009), but these must be removed post-production to restore antimicrobial activity to the AMP, adding an extra cost to production. A variety of ingenious methods have been proposed to perform the cleavage event without the use of proteases post-production (Tian *et al.*, 2009; Ke *et al.*, 2012), but, with one exception (Rothan *et al.*, 2014), AMPs that retain antimicrobial activity while still bound to the fusion partner have not been produced in bacteria. Plant expression of AMPs is an attractive alternative, since they are not themselves targeted by AMPs and have potential as highly scalable protein production systems. However, the yields so far reported for plant expression of AMPs (Lee *et al.*, 2010; Patino-Rodriguez *et al.*, 2013; Bundó *et. al.*, 2014) have been much lower than those reported for *E. coli* expression systems (Li, 2011). Even if production levels were competitive with *E. coli* systems, downstream processing contributes the bulk of production costs (Wilken and Nikolov, 2012), and this must be addressed especially for low-cost/large-scale applications.

We have expressed AMPs in a plant expression system and have addressed the two roadblocks mentioned above, achieving high expression of AMPs in plants and avoiding the carrier protein cleavage step, using a simple purification protocol. To increase yield of AMP fusion proteins in plants, we examined factors that might be responsible for low plant yield. We found that peptide charge was correlated with yield, as all of the anionic AMP fusions we tested were expressed in plants while none of the cationic peptides produced any detectable AMP fusion protein. To reduce downstream processing costs, we used an elastin-like polypeptide (ELP) carrier protein (Floss *et al.*, 2009), which confers to the fusion protein insolubility at 37°C, at which most protein contaminants are soluble, and solubility at 4°C. Centrifugation at 37°C pellets the fusion protein, which is then resuspended at 4°C. Unexpectedly, we found that the ELP-AMP fusions had antimicrobial activity without a protease cleavage step, which should further reduce post-production costs. This activity was, in fact, at least 10x stronger than that typically reported for cleaved AMPs produced in *E. coli* expression studies (Wei *et al.*, 2005; Tian *et al.*, 2009) or from AMPs synthetically produced (He and Lazaridis, 2013; Ebbensgaard *et al.*, 2015; Kubicek-Sutherland *et al*., 2017). Thus, this system fully leverages the potential unique advantages of plant production of AMPs as compared to other modes of production. The described method may thus serve as an antibiotic replacement platform for applications requiring large-scale, low-cost protocols.

## Materials and Methods

### Computation of hydrophobic ratio and net charge distributions from published sequences of STP-AMPs

AMP databases (PMID: 26602694, PMID: 18957441) were examined to determine correlations between taxonomic distribution and two protein structural factors, hydrophobicity and net charge. First, candidate peptide sequences were collected. To reduce other factors in the comparisons, the only AMPs examined were those having the most commonly occurring AMP structure, namely the sequential tri-disulfide peptide (STP) structure (Islam *et al.*, 2015; Islam *et al.*, 2018b). To collect STP-AMPSs, AMPs ranging from 30 to 50 amino acids were manually processed through the PredSTP tool (Islam *et al.* 2015) and the resulting peptide sequences were collected. We used the CD-HIT [PMID: 23060610] program to remove redundant sequences by setting a cutoff of sequence identity at 80%. The remaining sequences were grouped into the plant or non-plant origins using the original metadata.

After the candidate peptide sequences were collected, the hydrophobic ratio and the net charge of each sequence was calculated applying the identical formula used in the ADP3 server (PMID: 18957441). For the hydrophobicity calculation, A, I, L, M, W, V, C and F were considered hydrophobic amino acids, as shown below, with *n* being the number of occurrences of each corresponding amino acid in the peptide and *L* being the total number of amino acids in the peptide.

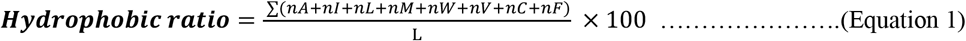

For the net charge calculations, the difference between the counts of negative (D + E) and positive (R + K) amino acids was defined as net charge for each peptide:

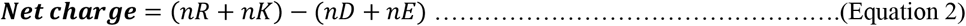

ANOVA tests were performed using R version 3.4.0 to observe any significant difference in the mean of hydrophobic ratio or net-charge in the STP-AMP sequences from plant or non-plant origins.

### Preliminary vector work

In preliminary experiments, two carrier proteins were tested as fusion partners for AMP transient expression in *N. benthamiana* via leaf agroinoculation. This work was performed before the meta-analysis of peptide charge described above. First, AMP was fused to the C-terminus of Jun a 3, a protein that expresses strongly and accumulates well in the apoplast of tobacco (Moehnke *et al.*, 2008). The Jun a 3 fusion was expressed using the plant viral vector FECT (Liu and Kearney, 2010). The AMPs tested were C16G2(+9), tachystatin B-1 (+7), protegrin (+6) and circulin-A(+2). No AMP expression was detected by SDS-PAGE/Coomassie Blue analysis (data not shown). Second, AMP was fused between eGFP and hydrophobin in the plant expression vector pCaMterX (Joensuu *et al.*, 2010). The AMPs tested were C16G2(+9), tachystatin B-1 (+7), sarcotoxin (+5), circulin-A(+2) and laterosporulin (−1). No AMP expression was detected by SDS-PAGE/Coomassie Blue analysis (data not shown). However, some GFP fluorescence was noted in plants inoculated with the anionic laterosporulin construct. In addition, the anionic insecticidal STP, Hv1a (−1), used as a positive control, expressed well in both of these systems. These were the first experimental data suggesting that peptide net charge may be a factor in the successful plant expression of AMPs.

### ELP vector

All subsequent work in the comparative expression of AMPs of different net charge was carried out using the ELP carrier protein. The ELP used in our study comprised 28 units of VPGVP pentapeptide repeats fused to the protein of interest (Conley *et al*., 2009b). The pCaMterX/ELP vector was modified to include unique restriction sites to allow insertion of AMP open reading frames (ORFs) with the excision of the eGFP ORF native to the original vector (Figure 1). Additionally, a TEV protease cleavage site (ENLYFQ) was inserted at the C-terminus of the AMP. The final construct allows for insertion/replacement at three sites for marker genes, AMPs and purifications tag (Figure 1).

**Fig. 1.**
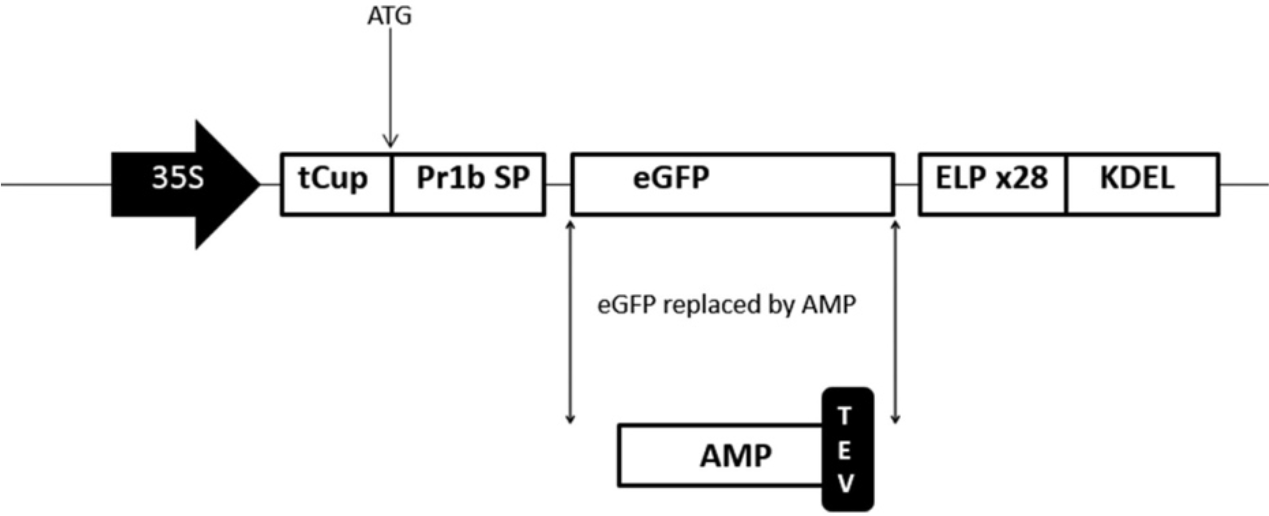
STP-AMP/Elastin-like polypeptide fusion sequence expressed via agroinoculation in *Nicotiana benthamiana*. 35S, CaMV 35S dual enhancer promoter; tCUP, translational enhancer; Pr1b SP, tobacco secretory signal peptide; KDEL, ER retention signal; TEV, tobacco etch virus protease recognition site (ENLYFQ).

### Agroinoculation, protein purification and analysis

*Agrobacterium tumefaciens* strain GV3101 was electroporated with the AMP/ELP binary vector and agroinoculation proceeded as described (Liu and Kearney, 2010), including the silencing suppressor, p19. All experiments were done in triplicate. As negative and positive controls, uninfected leaves and leaves infected with the original eGFP/ELP construct were collected and processed through the same ELP extraction and purification processing as the AMP/ELP samples.

AMP/ELP purification was performed as previously described (Conley *et al.*, 2009a). Specifically, plant leaves collected at 3-4 days post-inoculation were frozen in liquid nitrogen and ground with a pre-chilled mortar and pestle, then homogenized in three volumes (v/w) of ice cold 1X PBS. Extract was centrifuged in 4°C at 20,000 × g for 15 minutes. For the temperature-dependent inverse transition cycling, the supernatant above was warmed in a 37°C water bath with NaCl added to a concentration of 3 M. After 15-45 minutes of incubation extract was centrifuged at 37°C for 20,000 × g for 15 minutes. Supernatant was discarded, and the pellet was resuspended in ice cold 1X PBS at 1/10^th^ the volume and centrifuged at 4°C at 20,000 × g for 15 minutes. The resulting supernatant was the uncleaved protein product used for microbial inhibition studies.

To test the effect of cleavage on the fusion protein’s toxicity, AMP/ELP protein from the resuspended pellet was cleaved with TEV protease at a mass ratio of 4:1 in TEV protease buffer (50 mM tris HCl (pH 8.0), 0.5 mM EDTA, 1 mM DTT). The cleavage products (AMP and ELP) were not separately isolated and were analyzed as a mixed solution.

Protein extracts were analyzed by SDS-PAGE and mass spectrometry. Recombinant AMP protein yield was assessed by densitometry of SDS-PAGE band images measured against a BSA standard using NIH ImageJ. Mass spectrometry was used to confirm the presence of intact AMP and carrier peptide after TEV protease treatment and to confirm the identity of AMP-ELP fusion peptide from extracts not treated with TEV protease. Specifically, AMP/ELP fusion protein was first extracted from leaves using two cycles of the temperature shift protocol described above and TEV protease was used to cleave the fusion peptide into AMP and ELP. Cleaved or uncleaved fusion peptide was digested with trypsin and analyzed using LC-ESI-MS (Synapt G2-S, Waters) at the Baylor University Mass Spectrophotometry Center, followed by data analysis using MassLynx (v4.1). The results can be found in Supplementary Figures 5-40.

### Antibacterial assay of recombinant AMPs (MIC assays)

Purified AMPs and AMP/ELP fusion peptides were tested for their antimicrobial activity using a Minimum Inhibitory Concentration (MIC) assay. Specifically, 10 mL of *Staphylococcus epidermidis* was grown overnight in a shake culture at 150 rpm at 37°C. Turbidity was assessed with McFarland standard tubes and the culture was diluted to 0.5 OD_600_. The peptide was first added to the first well of a 96-well microtiter plate and serial 1:2 dilutions of the peptide were made across the plate using fresh LB medium. Then, 100 μl of *S. epidermidis* culture was added to each well containing the peptide dilutions and the culture was allowed to grow in the well at 37°C without shaking. To measure bacterial growth, resazurin was added to 0.00015% and plates were allowed to grow an additional 30-120 minutes until dye color changed to indicate bacterial growth or inactivity. All MIC experiments were run in triplicate

## Results

### STP-AMPs native to plants are less cationic than those from non-plant sources

We questioned why AMPs from non-plant sources generally express poorly as foreign genes in plant expression systems. We hypothesized that there may be a certain property characteristic to native plant AMPs that is not generally present in AMPs from other sources. We postulated that this property is necessary for robust expression of any AMP, from plant or non-plant origin, in heterologous plant expression systems. We selected AMP hydrophobicity and AMP net charge as two properties worth investigating.

We used a publicly available AMP database to access AMP sequences and metadata, but first applied filters to narrow the pool to those peptides of greatest practical value for heterologous expression in plants. Since we were most interested in peptides possessing the highly stable sequential tri-disulfide peptide (STP) structure, we used our PredSTP algorithm (Islam et al., 2015) to narrow the pool of AMPs gathered from the AMP database to only STPs. We further narrowed the pool to only peptides 30-50 amino acids in length and eliminated redundant sequences (80% sequence similarity cutoff), resulting in a final data set of 96 STP-AMPs of plant origin and 58 STP-AMPs of non-plant origin (Supplemental File 1).

Once the plant and non-plant STP-AMPs groups were collected, we compared them for hydrophobicity and net charge. We found no significant difference between the two groups in hydrophobicity (Figure 2). However, peptides of plant origin were found to be significantly less cationic than peptides of non-plant origin (Figure 2). A p-value of 4.47e-05 was determined by ANOVA for this comparison, with mean net charges of +1.77 versus +3.46 for STP-AMPs of plant vs. non-plant origin, respectively. Therefore, an unfavorably positive net charge may have been responsible for the poor expression of non-plant AMPs expressed in plant expression systems as reported in the literature to date.

**Fig. 2.**
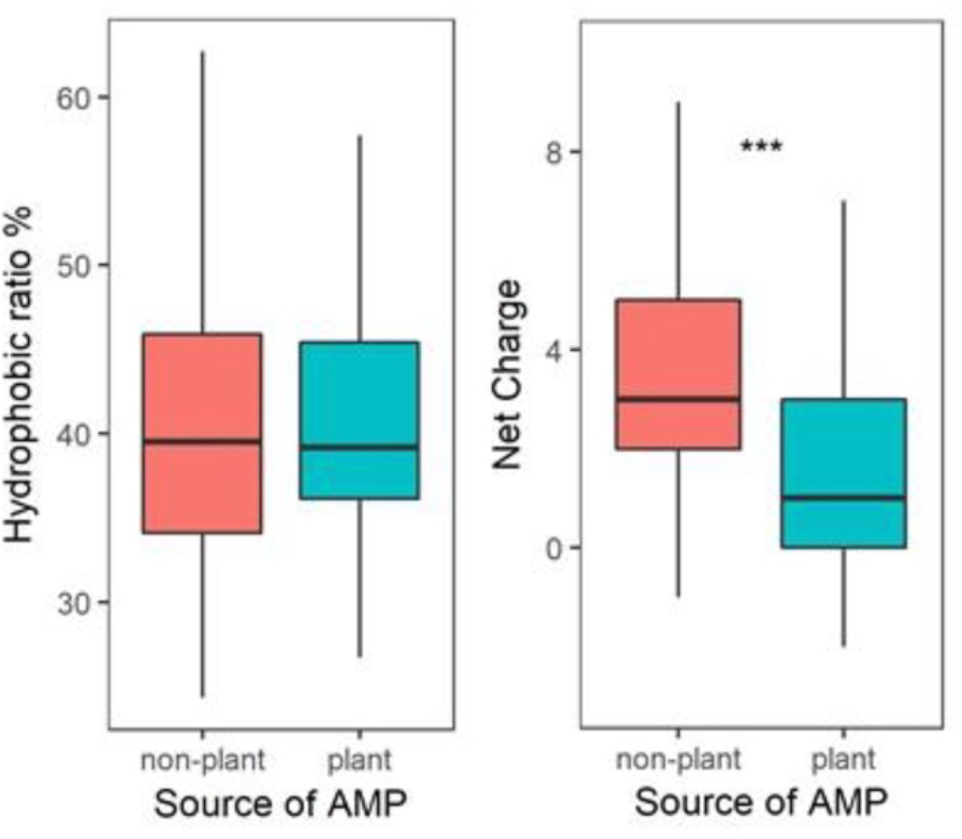
Meta-analysis of AMPs from Antimicrobial Peptide Database 2. Hydrophobicity and net charge were calculated and compared for STP-AMPs native to plant versus non-plant sources.

### Only anionic STP-ATP/ELP fusion proteins were expressed in transiently transgenic plants

From these findings, we formed the hypothesis that the expression of AMPs in plant expression systems may be improved by using AMPs which were anionic, neutral, or only slightly cationic. We tested this experimentally by expressing in the tobacco *Nicotiana benthamiana* a set of 10 AMPs ranging in net charge from highly cationic (+7) to highly anionic (−5). To eliminate the variables of peptide size, peptide structure and plant vs. non-plant origin, we selected only AMPs of 30-50 amino acids in length, possessing a core STP structure, and being of non-plant origin (Table 1).

**Table 1.**
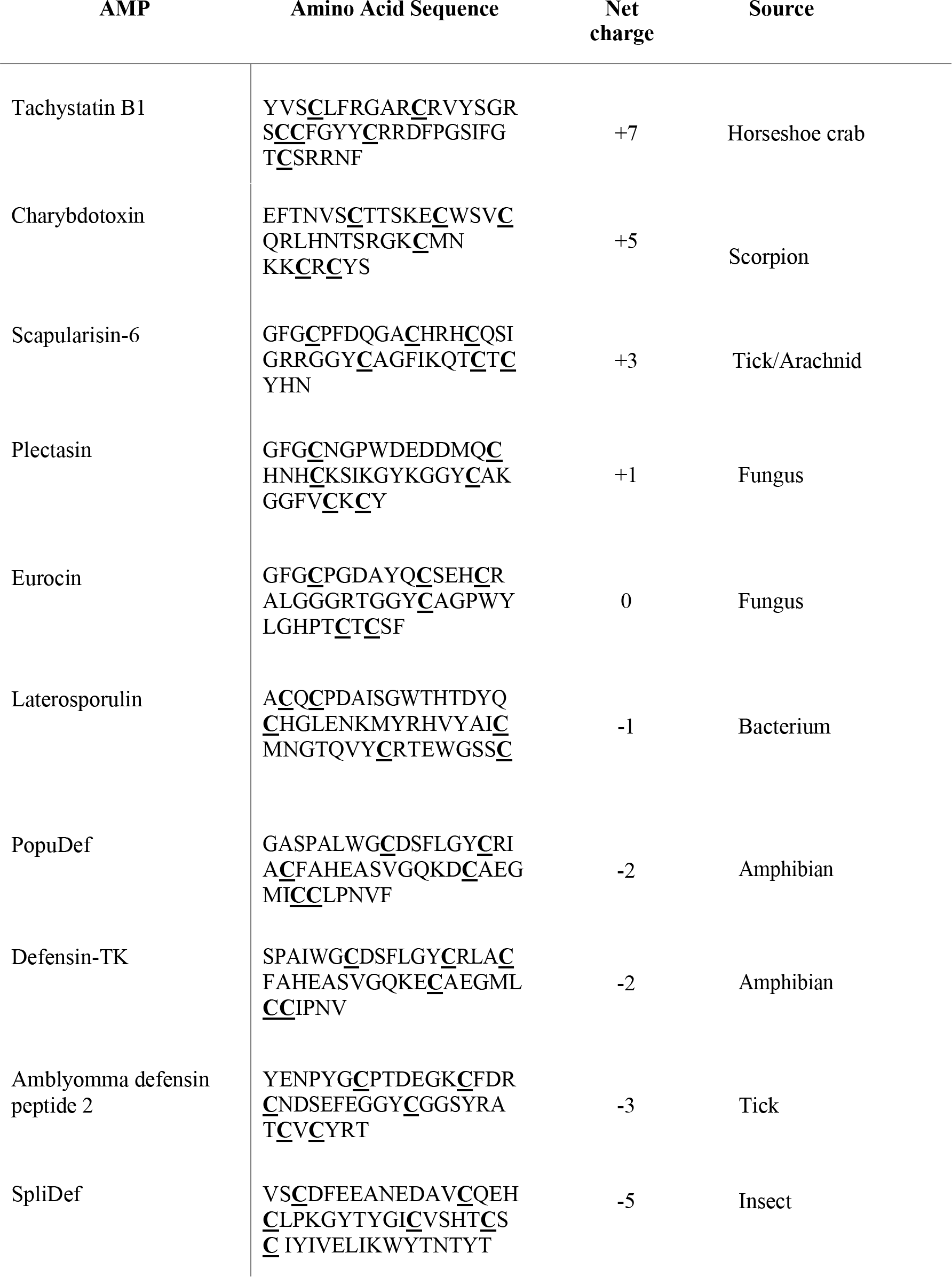
STP-AMPs cloned as ELP fusions and agroinoculated into *Nicotiana benthamiana*. The six cysteines participating in disulfide bonding in the STP structure are underlined.

When this range of 10 AMP/ELP fusions were expressed in *N. benthamiana* leaves, peptide net charge was seen linked to both yield and plant symptoms. Plants inoculated with cationic peptides showed a strong tendency to develop necrosis in the agroinoculated leaves and this effect was more severe the more cationic the peptide (Figure 3, top row). The neutral AMP, eurocin, and all anionic AMPs (bottom row) induced no leaf necrosis when agroinoculated as AMP/ELP fusion peptides. In line with these symptom observations, no expression of AMP/ELP was detected by SDS-PAGE with any of the cationic AMPs, nor with the neutral AMP, eurocin.

**Fig. 3.**
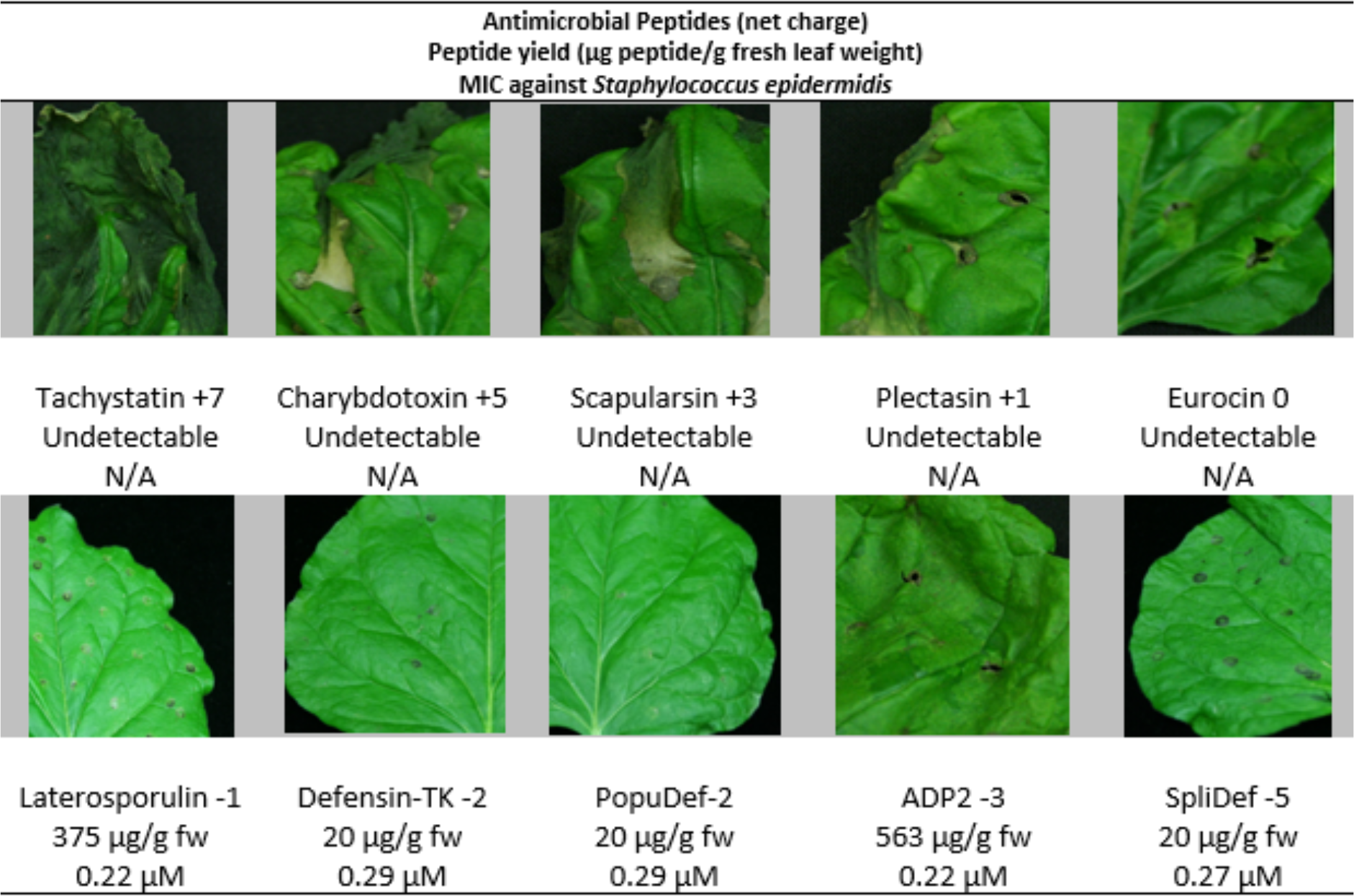
Expression of AMPs as ELP fusions in *Nicotiana benthamiana* via agroinoculation. Photos are displayed in order of AMP net charge, from most cationic to most anionic, showing a corresponding decrease in leaf necrosis. Peptide yield and minimum inhibitory concentration (MIC) values against *Staphylococcus epidermidis* for purified fusion peptides are indicated in the second and third line under each photo. The circular necrotic spots seen in all photos result from mechanical injury at agrobacterium injection points.

In contrast, every anionic AMP tested expressed as an AMP/ELP fusion to levels detectable by SDS-PAGE as a simple extract (Figure 4). The lowest levels of expression of anionic AMP/ELP fusions yielded 20 µg/gram fresh weight, which is comparable to the highest levels reported for plant expression of AMPs (refs). Furthermore, for two of the anionic AMP/ELP fusions, we noted over 10x greater expression with average yields of 375 and 563 μg/gram fresh weight for laterosporulin-1 and ADP2-3, respectively (Figure 3). This corresponds to 180 and 225 µg/gram fresh weight for each of the individual AMPs without the ELP carrier.

**Fig. 4.**
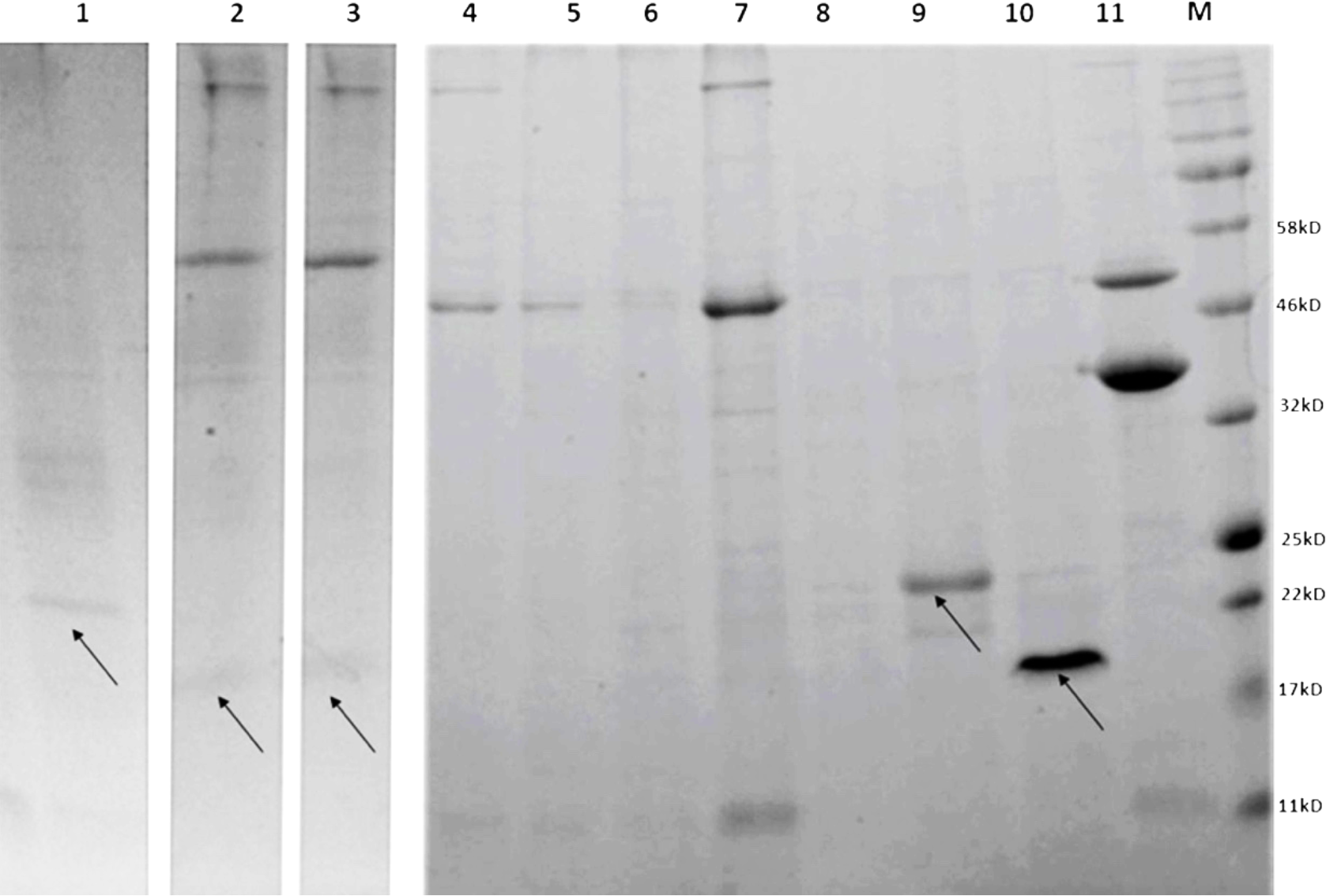
SDS-PAGE/Coomassie blue analysis of AMP/ELP fusion peptides expressed in *Nicotiana benthamiana*. AMP fusion protein expression was detected only for the anionic AMPs (arrows). Expression was especially strong for laterosporulin and ADP2 (Lanes 9 and 10). Lane 1, Defensin-TK; Lane 2, PopuDef; Lane 3, SpliDef; Lane 4, Tachystatin B1; Lane 5, Charybdotoxin; Lane 6, Scapularsin-6; Lane 7, Plectasin; Lane 8, Eurocin; Lane 9, Laterosporulin; Lane 10 ADP-2; Lane 11, ELP-EGFP positive control (35 kDa).

In SDS-PAGE analysis (Figure 4), extracts representing only one temperature shift cycle, with no further purification, were loaded onto the gel in order to demonstrate the purity of this relatively crude extract. The results also demonstrate the reliable yield obtained, as bands were clearly detectable for all anionic AMP/ELP peptides with standard Coomassie Blue staining even without any further concentration steps or nickel columns.

### Uncleaved AMP/ELP fusions had strong antibacterial activity

AMP/ELP fusion peptides of all of our anionic AMPs had unusually strong antibacterial activity as simple, unprocessed extracts. Against *Staphylococcus epidermidis*, our fusion peptides had MIC values that were consistently low (highly antibacterial), ranging from 0.22 - 0.29 μM for all AMP/ELP fusions (Figure 3). In contrast, the published MIC values against the related *Staphylococcus aureus* for the same AMPs purified from their source organisms are 7.5 µM for ADP-2 and 2 µM for laterosporulin (Lai *et al.* 2004; Singh *et al.*, 2012), which shows greater than 10-fold less antibacterial activity. Our GFP/ELP vector control gave an average MIC of 11.9 μM, demonstrating that the ELP carrier protein itself did not contribute significantly to antibacterial activity.

Attempts were made to find antibacterial activity in protease-treated extracts, but no activity was detected. To ensure that intact AMP was present after cleavage of the AMP/ELP fusion with TEV protease, protein analysis was performed by LC-EIS-MS for all fusions in the study that were successfully expressed in plants. For all of these, fully intact AMP was shown to be present in both the cleaved and uncleaved AMP/ELP fusion protein preparations (Supplementary Figures 5-40). Thus, strong antibacterial activity was demonstrated in the uncleaved AMP/ELP fusion proteins but no activity was found in the protease-treated extracts, despite the presence of intact AMP.

## Discussion

As a protein expression system, plants bring the unique potential advantage of low-cost production and scalability. However, the yields of antimicrobial peptides reported from plant systems to date are far lower than those from *E. coli* and other competing expression systems, suggesting an intrinsic incompatibility between the plant hosts and the heterologous AMPs expressed. This presents a barrier to commercialization, with yields insufficient to take advantage of the scalability of plant systems. As an example, the synthetic cationic AMP BP100 showed phytotoxicity in *Arabidopsis* seedlings and fitness reduction in rice plants (Nadal *et al.*, 2012), and had relatively low yield in *N. benthamiana* leaves (Company *et al.*, 2013). As another example, seed expression systems often provide high yields and the expression of the AMP cecropin A in rice seed endosperm did not negatively impact seed physiology. Even so, the yield was low, ranging from 0.5-6 µg per gram seed tissue weight (Bundó *et. al.*, 2014). Taking another approach, protegrin-1 (PG1) was expressed in the powerful magnICON tobacco mosaic virus (TMV) vector, with the AMP directed to the apoplast of *N. tabacum* leaves, but no yield figures were reported. Another powerful expression system involves chloroplast expression, which has the added advantage of being prokaryotic in nature. However, when protegrin was produced as a fusion with GFP in a chloroplast expression vector, the yield of the purified fusion protein was only 8 µg/g fresh weight of leaf tissue (Patino-Rodriguez *et al.*, 2013; Lee *et al.*, 2010). Finally, fusing AMPs to carrier proteins is normal practice in *E. coli* expression of AMPs and this was attempted with sarcotoxin IA, using GUS as the carrier protein for plant expression. However, the levels expressed were not sufficient for detection by SDS-PAGE (Okamoto *et al*., 1998).

In our study, we appear to have broken the yield barrier for AMP expression in plants by observing a bias in peptide charge found naturally in plants and then experimentally demonstrating that, for our set of 10 AMPs, only the anionic AMPs could be expressed. Our minimum yields for anionic AMPs were slightly above the highest reported AMP yields in plant expression systems to date; furthermore, our highest yielding AMPs delivered 10-fold as much. These yields may be compared to those in *E. coli*, which, typically, produces 10-100 mg of AMP from a 1-liter culture (Li, 2011). In comparison, reported yields from the previously published plant expression systems cited above would correspond to 1 mg from a medium-sized harvest of 200 g of plant tissue. In contrast, we report an AMP yield in plants which is comparable to that achieved in *E. coli*. In our study, a minimum yield of 20 μg and a maximum of 563 μg per gram fresh weight was observed, which would correspond to 4 mg and 113 mg per 200 g of plant tissue, respectively, equivalent to reported yields for *E. coli* expression systems. In perspective, the best yields of anionic AMP/ELP fusion peptides in our study also compare favorably to reports for the expression of the marker gene GFP in *N. benthamiana* plants (270-340 μg GFP/gfw) using a 35S promoter aided by the p19 silencing suppressor (Voinnet et al., 2003).

Furthermore, the AMP/ELP fusion peptides of our study possess an antimicrobial activity (0.22-0.29 µM) an order of magnitude stronger than these AMPs expressed from *E. coli* systems (Li *et al.*, 2010; Parachin *et al.*, 2012; Li *et al*., 2017; Mao *et al*., 2013). Thus, on a functional basis, the yield figures we report would be considerably higher in comparison to those of the *E. coli* systems.

The use of elastin-like polypeptide (ELP) as a fusion partner proved important for the yield and antibacterial activity of anionic AMPs expressed in plants in this study. ELP is an extracellular matrix protein found in vertebrate connective tissue. When targeted to the endoplasmic reticulum, ELP provides protein sequestering and stability to its fusion partner (Floss *et al.*, 2009; Sousa *et al*., 2016; Streatfield *et al.*, 2007). The ELP protein also provides a purification process using inexpensive temperature shifts without the use of chemicals or chromatography (Floss *et al*., 2009; Meyer and Chilkoti, 1999). In addition to benefiting yield and purification, we noted that the unusually high antibacterial activity was associated with the uncleaved ELP fusions. Activity of uncleaved ELP/AMP fusions has not been previously reported in *E. coli* expression systems. Further studies are in progress in our lab to elucidate the protein structural aspects of antimicrobial activity of AMP fused to the ELP carrier.

Plant expression of AMPs seems well suited to large scale, low-margin applications and the effectiveness of AMPs has already demonstrated as poultry and livestock feed additives (Juarez *et al.*, 2016), food preservatives (Rai *et al.*, 2016) and topical disinfectants (Pfalzgraff *et al.*, 2018). The scalability of plant expression systems would allow for the production of large amounts of raw product, which could then be reduced to relatively pure protein by simple temperature shift cycles, potentially without the need for column chromatography, which increases post-production costs (Wilkin and Nikolov, 2012). Alternatively, the AMPs might be expressed in transgenic grain seed, which tends to have yields higher than seen in leaf tissue. Recombinant proteins remain stable using traditional seed storage technique (Boothe *et al.*, 2010; Morandini *et al*., 2011; Hnatuszko-Konka *et al.*, 2016). Anionic AMPs used in the seed platform would address the increasing concern over the amount of antibiotics used with livestock (Ferber, 2003; Massé *et al.* 2014; Hao *et al*., 2015). Applications in animal feed have already shown limited success in protecting livestock against pathogens using antibodies and antimicrobial peptides (Virdi *et al*., 2013; Lee *et al.*, 2010). Since the ELP/AMP fusion protein does not need to be cleaved, it would be expected that the resulting grain should be directly antimicrobial, without proteolytic processing. Proteins encapsulated in plant tissue may be expected to better survive the digestive system to arrive intact to eliminate gut pathogens (Sabalza *et al.*, 2013). Furthermore, AMPs would largely eliminate the environmental consequences of pesticide use. Being peptides, AMPs would be expected to have a very short half-life in soil or aquatic environments.

A final advantage to using AMPs is to avoid the development of resistance in target bacteria and the microbiota as a whole (Perron *et al.*, 2006; Maróti *et al*. 2011). As already mentioned, amphipathic AMPs have a generalized mechanism for destroying bacteria by membrane disruption and other mechanisms, which inhibits resistance development (Peschel and Sahl, 2006; Nguyen *et al.*, 2011). Furthermore, it would be relatively easy to supply AMPs as "stacked drugs" by including several AMPs in the same treatment or in the same grain seed genome. In this way, resistance development is further forestalled since any resistance mutant that appears would be destroyed by another AMP with a divergent mode of action in the treatment mix. The sheer abundance of putative AMP sequences in genome databases is a vast resource for the rapid discovery and testing of large numbers of AMPs to support a stacked drug paradigm, as opposed to the slow development cycle of small molecule antibiotic drugs. We have added to the capacity of this developmental pipeline by developing several algorithms for detecting AMPs from genomic databases. We can now predict sequential tri-disulfide peptides (STPs) from genomes using a support vector machine algorithm (Islam *et al.*, 2015). STPs are the predominant structural form of AMPs and are study, robust structures (Islam *et al.*, 2018b). We have developed a generalized algorithm based on natural language processing to classify protein sequence based on any input characteristic (Islam *et al.*, 2018a) and this has been used to predict protein function from genome sequences, such as picking AMP function while rejecting hemolytic activity to avoid human toxicity (Islam *et al.*, 2018c). Anionic AMP candidate sequences from these genome searches can be expressed as ELP/AMP fusion peptides in plants with the expectation that a significantly proportion would express at high yield, as we observed in our present study. These fusion peptides could then be screened as extracts from a single thermal shift cycle, accelerating the screening workflow process.

## Acknowledgements

We thank the Baylor Mass Spectrometry Center and Molecular Biology Center for expert advice and facilities.

